# Foot form and function: Variable and versatile, yet sufficiently related to predict one from the other

**DOI:** 10.1101/2022.10.02.510569

**Authors:** Robert W. Schuster, Andrew Cresswell, Luke A. Kelly

**Affiliations:** School of Human Movement & Nutrition Sciences, The University of Queensland, Australia

**Keywords:** foot morphology, foot mechanics, longitudinal arch, transverse arch, shape-function modelling

## Abstract

The modern human foot is a complex structure thought to play an important role in our ability to walk and run efficiently. Comparisons of our feet to those of our evolutionary ancestors and closest living relatives have linked the shape of several foot components (e.g., the longitudinal and transverse arches, size of the heel and length of the toes) to specific mechanical functions. But since foot shape varies widely across the modern human population, this study aimed to investigate how closely foot shape, deformation and joint mechanics during various locomotor tasks are actually linked. And whether the latter can be accurately predicted based entirely on the former two. A statistical shape-function model (SFM) was constructed by performing a principal component analysis on 100 participants’ three-dimensional foot scans, and joint angles and moments captured during level, uphill, and downhill walking and running. This SFM revealed that the main sources of variation were the longitudinal and transverse arches, relative foot proportions and toe shape along with their associated joint mechanics. However, each of these only accounted for a small proportion of the overall variation in foot shape, deformation and joint mechanics, most likely due to the high structural complexity and variability of the foot. Nevertheless, a leave-one-out analysis showed that the SFM can be used to accurately predict the joint angles and moments of a new foot based only on its shape. These results have implications and potential applicability across numerous fields, such as evolutionary anthropology, podiatry, orthopaedics and footwear design.

## Introduction

The observation that the form and function of a biological system seem to be closely linked is a central aspect of the various theories of evolution. This idea is fundamental to Lamarck’s inheritance of acquired traits, Darwin’s survival of the fittest, as well as the more recent modern synthesis. Nature provides an astonishingly wide range of examples that support this form-function relationship; from the giraffe’s long neck that allows it to eat hard-to-reach leaves, the fennec fox’s large ears with which it can dissipate heat, all the way to the aye-aye’s unusual third digit for foraging larvae in hollow trees. And since modern humans evolved through the same (or at least similar) mechanisms as those that produced these highly specialised anatomical adaptations, it is only logical to assume that the unique features of our anatomy also serve a specific functional purpose. One unique functional ability, for which investigators have tried to identify the necessary anatomical adaptations, is the effectiveness with which we can walk and run in an upright position. A common method used in this endeavour, is comparing ourselves to our evolutionary ancestors and closest living relatives. These comparisons have produced a comprehensive list, ranging from our relatively flat face, unique spinal curvature, short and narrow pelvis, large hip and knee joints, long legs and short arms to the unique shape and structure of our feet (Bramble and Lieberman, 2004; Hunt, 2015).

Specifically, the last of this list, our feet, have received much attention. Despite their significance as our most common interface with the ground, the highly complex structure of our feet has made deciphering the relationship between their form and function far less straightforward than that of, for example, a giraffe’s neck. Consequently, a wide range of form-function relationships concerning the modern human foot have been proposed. Early discussions regarding the functional advantages our feet provide singled out their arched configuration; in particular the longitudinal arch (LA). This LA was recognized as characteristic of modern humans and thought to be a reflection of the forces transmitted and leverage provided by the foot (Morton, 1924). Attributed to the shape of the tarsal and metatarsal bones, and their joints, the LA is thought to be a source of the midfoot stiffness that makes our feet an effective lever to push off from when walking and running (Elftman and Manter, 1935; Hicks, 1955). However, the notion of LA stiffness has been challenged by investigations that have shown the midfoot to display considerable mobility (Holowka et al., 2017; Kessler et al., 2019; Lundgren et al., 2008; Welte et al., 2021; Welte et al., 2018). Through this mobility, the LA is also thought to increase locomotor economy by acting like a spring that stores and returns elastic energy with each step (Ker et al., 1987). Given the debate around LA stiffness and mobility, the transverse arch (TA) of the foot has been proposed as an alternative source of significant longitudinal stiffness (Venkadesan et al., 2020).Outside of the foot arches, the enlarged calcaneal tuber in humans compared to predominantly quadrupedal African apes is seen as an adaptation to the larger forces encountered when striking the ground during bipedal gait (Latimer and Lovejoy, 1989). And at the other end of the foot, the shorter toes in humans are considered an adaptation to increase running economy (Rolian et al., 2009).

Comparative studies such as these are generally based on a small number of feet assumed to be representative of an entire species. However, human foot form has been extensively documented to exhibit great variability (Cavanagh and Rodgers, 1987; Domjanic et al., 2015; Stanković et al., 2018), which, in turn, has led to the question whether variations in form are also related to variations in function *within* our species. Similar to the inter-species comparisons, several of the studies attempting to answer this question have focused on the LA. The shape of which has commonly been represented by either the height or change in height of the LA, while a far less constrained approach has been taken in regard to function, using measures such as lower limb kinematics (Nigg et al., 1993; Williams III et al., 2001), ground reaction forces (GRFs) (Nachbauer and Nigg, 1992; Williams III et al., 2014), and injury patterns (Hagedorn et al., 2013; Riskowski et al., 2013; Tong and Kong, 2013). This approach has led to conflicting results and a lack of general consensus, with some studies claiming LA shape to be an important predictor of how someone walks and runs or the injuries they are at risk of sustaining (Tong and Kong, 2013; Williams III et al., 2001), and others claiming the contrary (Nachbauer and Nigg, 1992; Nigg et al., 1993). Taking a more focused approach, DeSilva et al. (2015) found that LA height is related to midfoot mobility and midfoot peak plantar pressure during walking. Going into even more detail, the same investigation also showed that midfoot plantar pressure was also correlated to the curvature of the base of the fourth metatarsal. Further investigations regarding the shape or configuration of the bones in our feet have found that variations in calcaneus length can explain up to 80% of the variance in the metabolic cost of running (Raichlen et al., 2011) but that only 35% of the variance in dynamic plantar pressure can be explained by radiographic measures of foot structure (Cavanagh et al., 1997).

All of these previous investigations have used specific components of the foot to represent its form (e.g., skeletal structure, footprints, LA height). However, when applied individually these approaches might overlook important information that, if included, might help shed further light onto the relationship between the many variations in the shape of our feet and their mechanical function. Therefore, this study aims to determine how closely foot form and function are linked when representing the foot as an integrated whole by constructing a statistical shape-function model (SFM) from three-dimensional (3D) foot scans and the joint angles and moments measured during various locomotor tasks. Furthermore, this study also aims to evaluate the accuracy with which the mechanical function of a given foot can be predicted based solely on features of its external shape.

## Methods

### Participants

One hundred healthy adults, aged between 18 and 40 years, (50 female; 25 ± 6 years; 175 ± 9 cm; 72 ± 12 kg) were recruited to participate in this study. Individuals that had sustained a lower limb injury 12 months prior, or had any known neurological impairment, musculoskeletal or cardiovascular conditions were excluded from participating. Approval for the study protocol was attained from The University of Queensland’s institutional human research ethics committee (2019000999). All participants provided written informed consent prior to participation.

### Experimental Protocol

#### 3D Foot Scanning

To be able to quantify the deformations that occur when a foot is encumbered with increased load, the participants’ dominant foot was scanned (∼30 s duration) while bearing minimal load (mBW) and full bodyweight (fBW). Following is a brief description of the load application and scanning procedures as these are described in detail elsewhere (Schuster et al., 2021). For mBW, the load on the foot was minimized by having the participant seated. The seat height was raised to just below the height where the participant’s heel lost contact with the scanner’s surface. For fBW, participants were asked to stand upright, supporting a barbell equal to their body mass across their shoulders. Participants were then asked to distribute the combined weight of body mass and barbell equally between both feet. All scans were captured using a FootIn3D scanner (Elivision, Lithuania; reported accuracy 0.3 mm) with a 1.4 mm mesh resolution.

#### Treadmill Walking and Running

Through the combined efforts of its passive and active components, the foot is capable of transitioning between compliant spring, energy producing motor and energy absorbing damper during tasks with different work requirements at the centre of mass (Birch et al., 2021; Riddick et al., 2019; Smith et al., 2021). To determine how feet with different morphologies behave during such different tasks, participants were asked to walk and run (Froude number = 0.25 and 1.00, respectively) at three inclinations each (walking: -20, 0, 20%; running: -10, 0, 10%) on a force-sensing, fore-aft, tandem treadmill (AMTI, USA). Three trials were performed for each condition, during each of which a 12 s recording of GRFs and body segment kinematics was captured. GRFs were recorded at 1000 Hz while kinematics were synchronously recorded using a 14-camera 3D motion capture system (Qualisys, Sweden) sampling at 200 Hz. A total of 56 reflective markers (9.0 mm diameter) were used to represent the thigh, shank, rearfoot, midfoot, forefoot and toes of both legs and feet, as well as the pelvis. The marker placement defined by the Istituto Ortopedico Rizzoli foot model (Leardini et al., 2007) was used to represent four foot segments, with additional markers placed on the proximal interphalangeal joints of the second and fourth toes. Markers placed over the anterior and posterior superior iliac crests were used to represent the pelvis. The thighs and shanks were represented by markers placed over the lateral and medial malleoli and femoral epicondyles. Additional clusters of four markers were placed laterally on both shanks and thighs to more accurately track the movements of these segments.

### Data Processing and Analysis

#### Foot Shape and Deformation

A necessary precursor to building a statistical shape model is ensuring that all elements to be included in the model consist of the same number and order of vertices, and that they are spatially aligned. For this registration process an algorithm consisting of an iterative affine transformation (for anatomical alignment) and elastic deformation (for shape correspondence) was used (Danckaers et al., 2014). The scan with the closest surface area to the mean was chosen as the reference and registered to all other scans. The resulting group of uniquely shaped foot scans with 29528 corresponding vertices was further translated, rotated and scaled onto the reference through a General Procrustes Analysis. The 3D cartesian coordinates of the vertices from the registered and scaled fBW foot scans were then used as input for the SFM.

All scans, regardless of the load they were captured under, were registered and aligned to the same reference. Thus, Euclidean vectors between corresponding vertices on scans of the same foot captured under two loads can be used to describe the deformations that foot underwent due of an increase in load (Schuster et al., 2021). As with the vertex coordinates describing foot shapes, the components of the 29528 Euclidean vectors between mBW and fBW foot scans were used as the foot deformation input for the SFM.

#### Foot Kinematics and Kinetics

A 15 Hz low-pass, recursive second-order Butterworth filter was applied to all kinematic and kinetic data. The GRFs were first separated into periods of left and right ground contacts based on vertical GRFs, centre of pressure and foot marker positions. For each of these ground contacts, relative proportions of the GRFs were assigned to the three foot segments that contact the ground (calcaneus, metatarsals and toes) using a weighted probability algorithm described by Riddick et al. (2019). The marker positions from a static trial, captured prior to the walking and running trials, were used to scale an anatomically articulated multi-segment foot and ankle model (Maharaj et al., 2021) to each individual’s dimensions in OpenSim (Delp et al., 2007). Once scaled, the model was combined with the GRFs for inverse kinematic and dynamic analyses to determine ankle, subtalar, midtarsal, tarsometatarsal and metatarsophalangeal (MTP) joint motions and moments. The resulting joint angles and moments were time-normalised to 101 points for each stance phase and averaged across all strides during each locomotor task for each participant. These time-normalised and stride averaged joint angles and moments were used as the function input for the SFM.

### Statistical Analyses

To ensure a complete set of training data from which to build the SFM, joint angles and moments from missing uphill and downhill running trials were imputed for two participants. As recommended by Dray and Josse (2015), an iterative PCA algorithm, implemented in a specialised R-package (Josse and Husson, 2016), was used.

Following the methods described by Smoger et al. (2015), the SFM was constructed using the input variables for foot shape, deformation and function detailed above. Fig. 1 briefly describes this SFM construction process as well as how the resulting eigenvector matrix and principal component (PC) scores can be used to reconstruct the input data. A detailed description can be found in the supplementary information. The input data reconstruction was used to visualise the changes in shape, deformation and function described by the first five PCs, and to determine the accuracy of the SFM when predicting function from just shape or shape and deformation data. For the visualisations, the mean scores of each PC were perturbed by ± 3 standard deviations (SD) and then used to reconstruct the input variables. For the foot function prediction accuracy, a leave-one-out (LOO) analysis was performed, iteratively constructing the SFM while omitting one participant (***L***) from the standardised input data matrix (***Y***). Similar to the approach described by Fitzpatrick et al. (2011b), the standardised shape data from the left-out participant (***L***_***s***_) and the shape portion of the eigenvector matrix (***E***_***s***_) were used to determine shape PC scores (***S***_***Ls***_ = ***L***_***s***_***E***_***s***_). The same was done for shape PC scores of the training data (***S***_***s***_ = ***Y***_***s***_***E***_***s***_), after which the correlation coefficients (***r***) between these sample shape PC scores and the sample overall PC scores (***S*** = ***YE***) were determined. The overall PC scores for the left-out participant were then estimated using the correlation coefficients and left-out shape PC scores (***S***_***L***_ = ***S***_***Ls***_***r***). Using only the estimated overall PC scores from PCs with a corresponding ***r*** ≥ 0.95, the overall input data was reconstructed (***L’***_***raw***_ = ***S***_***L***_***E***^***T***^ ***σ*** + *μ*) and compared to the true shape, deformation and function data (***L***_***raw***_).

**Fig. 1.**
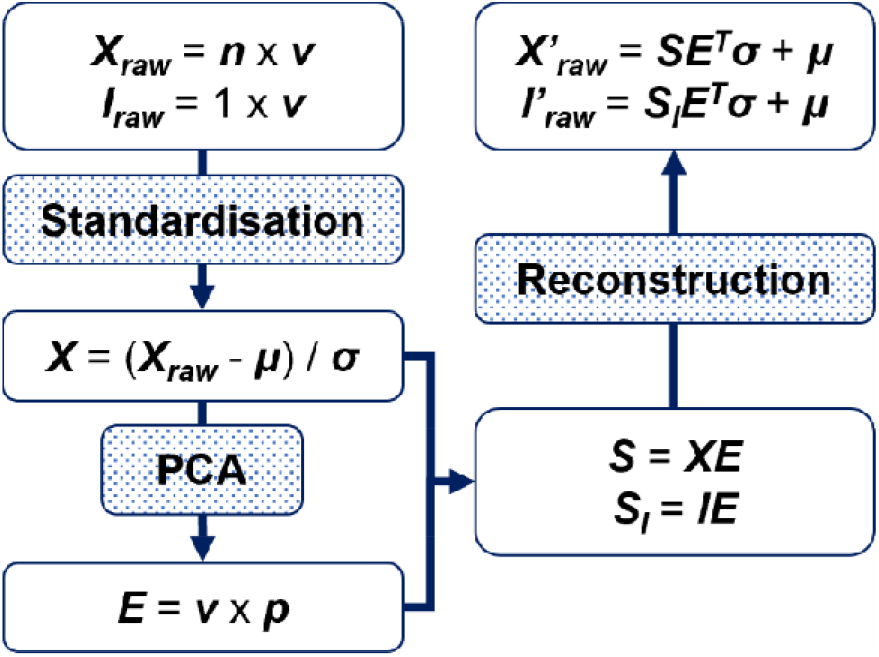
Schematic of the shape-function model construction process. Principal component analysis (PCA) is applied to the standardised input data matrix (**X**) consisting of **n** participants and **v** variables. The transpose of the resulting eigenvector matrix (**E**^**T**^), consisting of **v** variables and **p** principal components, can then be used together with the corresponding principal component scores (**S**) and group means (μ) and standard deviations (σ) to reconstruct the input data for all (**X’**_**raw**_) or a single participant (**I’**_**raw**_).

The accuracy of the shape and deformation reconstructions are reported as the sample mean ± SD of the absolute geometric error averaged across all vertices or vectors. The accuracy of predicted foot function is reported as the sample mean of the absolute error averaged across the entire stance phase for each joint, during each locomotor task.

## Results

### Statistical Foot Shape-Function Model

The SFM reduced the 183228 foot shape, deformation and function variables to 100 PCs. The first six of which captured 50.8% of the variability contained within the entire set of training data, while 16, 25 and 50 PCs explained 76.2%, 85.2% and 95.0%, respectively. These PCs also described the relationships between foot form and function identified by the SFM. To help understand these relationships, the changes in shape, deformation and function resulting from perturbations to these PCs are presented in Figs 3 - 7, as well as in figures and animations in the supplementary information. Since only the first five PCs accounted for more than 5% of the variability each, they will be the only PCs discussed in detail here.

**Fig. 2.**
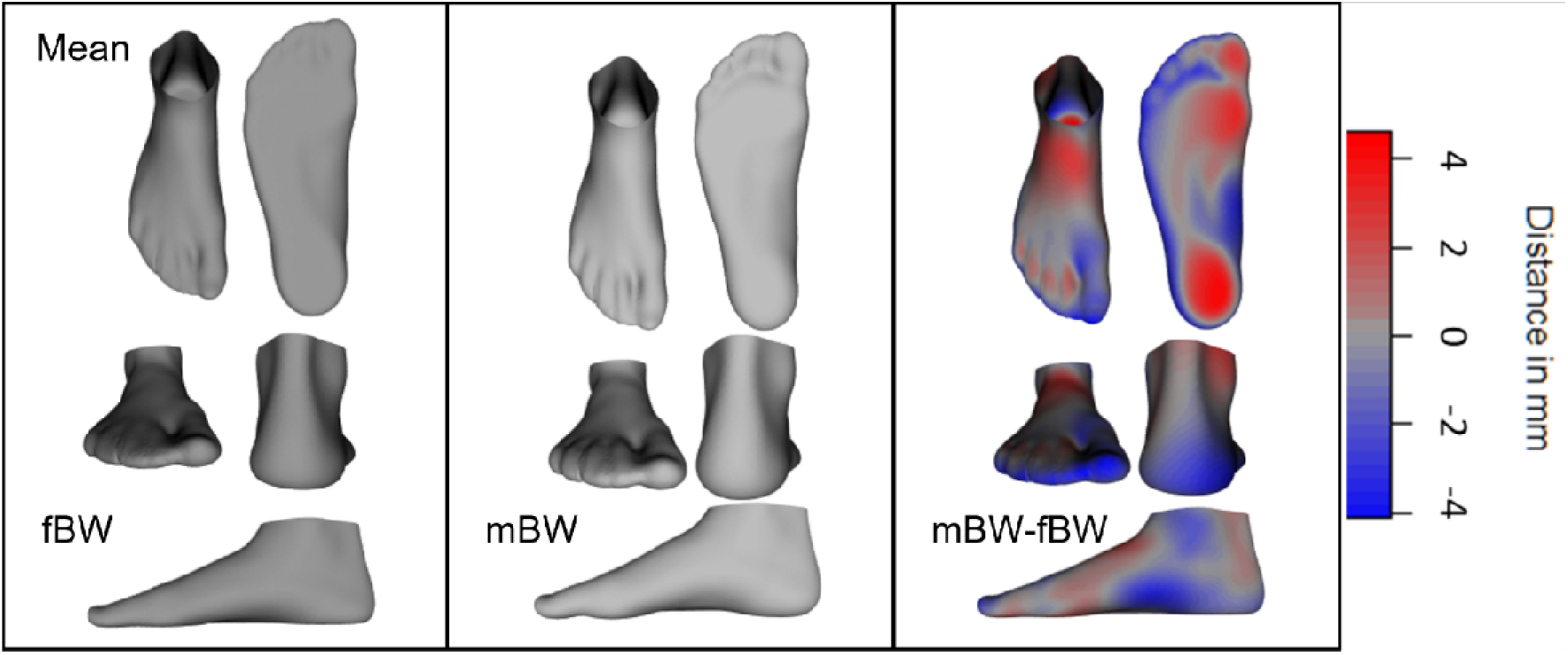
Mean foot shape while bearing full bodyweight (fBW) and minimal weight (mBW), as well as mean deformations when increasing the load from mBW to fBW (mBW-fBW). The colour coding on the mean deformed foot depicts the areas and magnitude of differences between the mean mBW foot and the mean deformed foot.

**Fig. 3.**
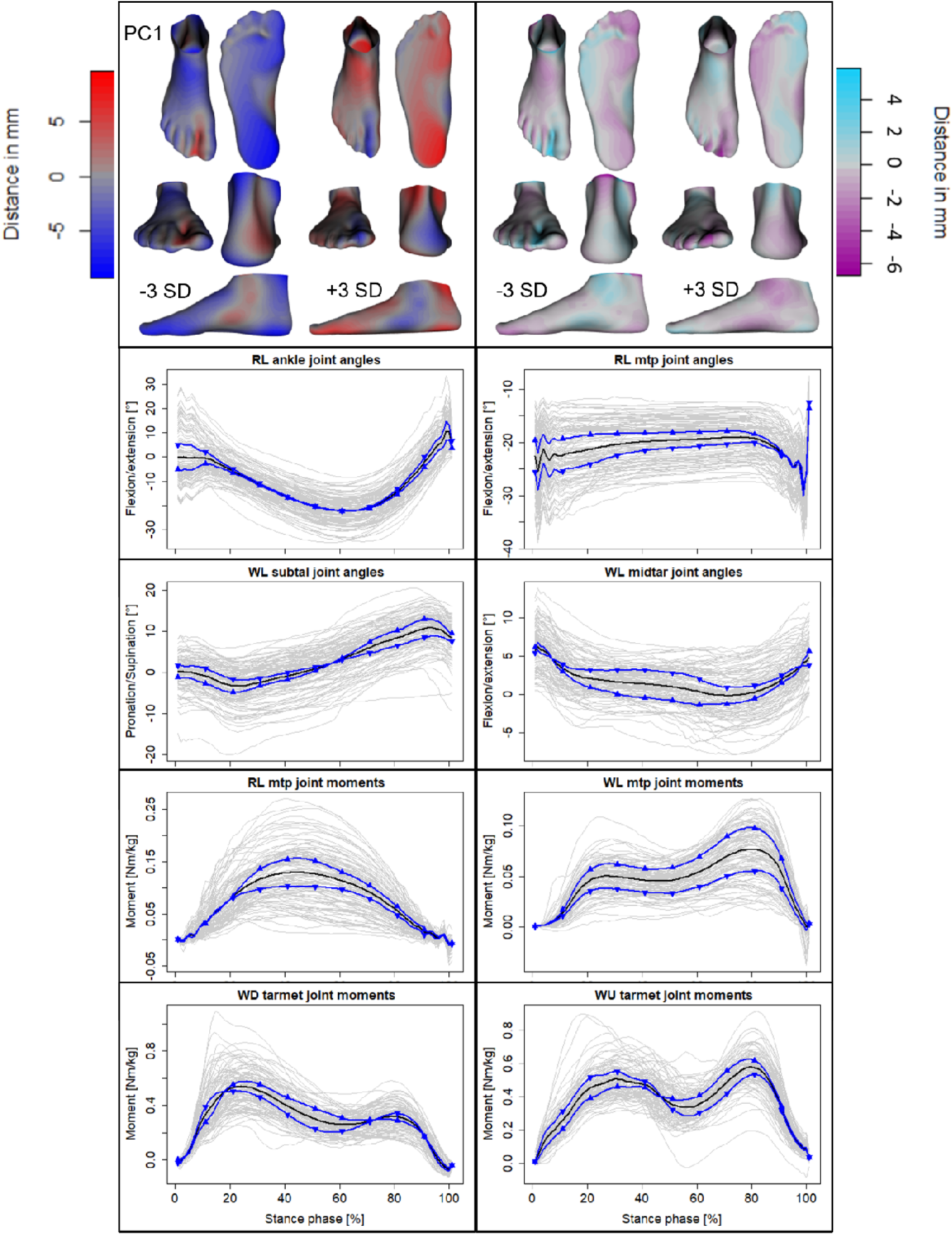
Mean ± 3 standard deviations (SD) of shape (top left), deformation (top right) and joint angle and moment variations described by the first principal component (PC). Colour coding depicts the degree of deviations from the mean foot or mean deformed foot (Fig. 2). Ankle, metatarsophalangeal (mtp), subtalar (subtal) and midtarsal (midtar) joint angles during level running (RL) and walking (WL) are shown. Joint moments of the mtp joint during RL and WL, as well as of the tarsometatarsal (tarmet) joint during downhill (WD) and uphill walking (WU) are shown. Upward pointing triangles (▲) denote +3SD and downward pointing triangles (▼) -3SD. Mean (black line) and participant means (grey lines) are also included as a reference.

The first PC explained 16.1% of the total variability and captured similar shape variations previously described by the first PC of statistical models concerned only with foot shape (Schuster et al., 2021; Stanković et al., 2018). These shape variations include differences in LA height, heel and forefoot width and toe splay. Associated with these shape characteristics were differences in LA compression, ankle internal/external rotation, and medial/lateral plantar compression as a consequence of increased loading. Specifically, supinated feet with a high LA and wide forefoot (i.e., -3 SD) exhibited ankle external rotation and limited LA compression but with substantial forefoot splay. These feet also appeared to contact the ground with the ankle plantarflexed or close to neutral instead of dorsiflexed, the MTP joints in a less dorsiflexed position, and the subtalar joint in supination instead of pronation, especially during downhill walking and all running tasks. Furthermore, they appeared to cover a smaller midtarsal joint range of motion (ROM) throughout stance than their pronated, low-arched counterparts (i.e., +3 SD), while exhibiting greater MTP joint dorsiflexion throughout the first 80% of stance and lower moments during midstance (∼20-80%). During uphill walking, high-arched feet exhibited greater ankle, subtalar, midtarsal and tarsometatarsal joint moments than low-arched feet during early stance and vice versa during late stance. When walking downhill, high-arched feet were linked to greater joint moments in late stance and with a steeper rise in early stance (Fig. 3).

The second PC accounted for 10.6% of the total variability. The shape differences contained within this PC primarily describe differences in the width of the midfoot, size and angle of the hallux, dorsal curvature as well as the relative lengths of the forefoot and heel. According to this PC, feet with a large, straight hallux, narrow midfoot and short heel (−3 SD) experienced greater rearfoot abduction, and less toe extension when loaded. They also seemed to exhibit larger ankle, subtalar, midtarsal, and tarsometatarsal joint moments during level running (Fig. 4).

**Fig. 4.**
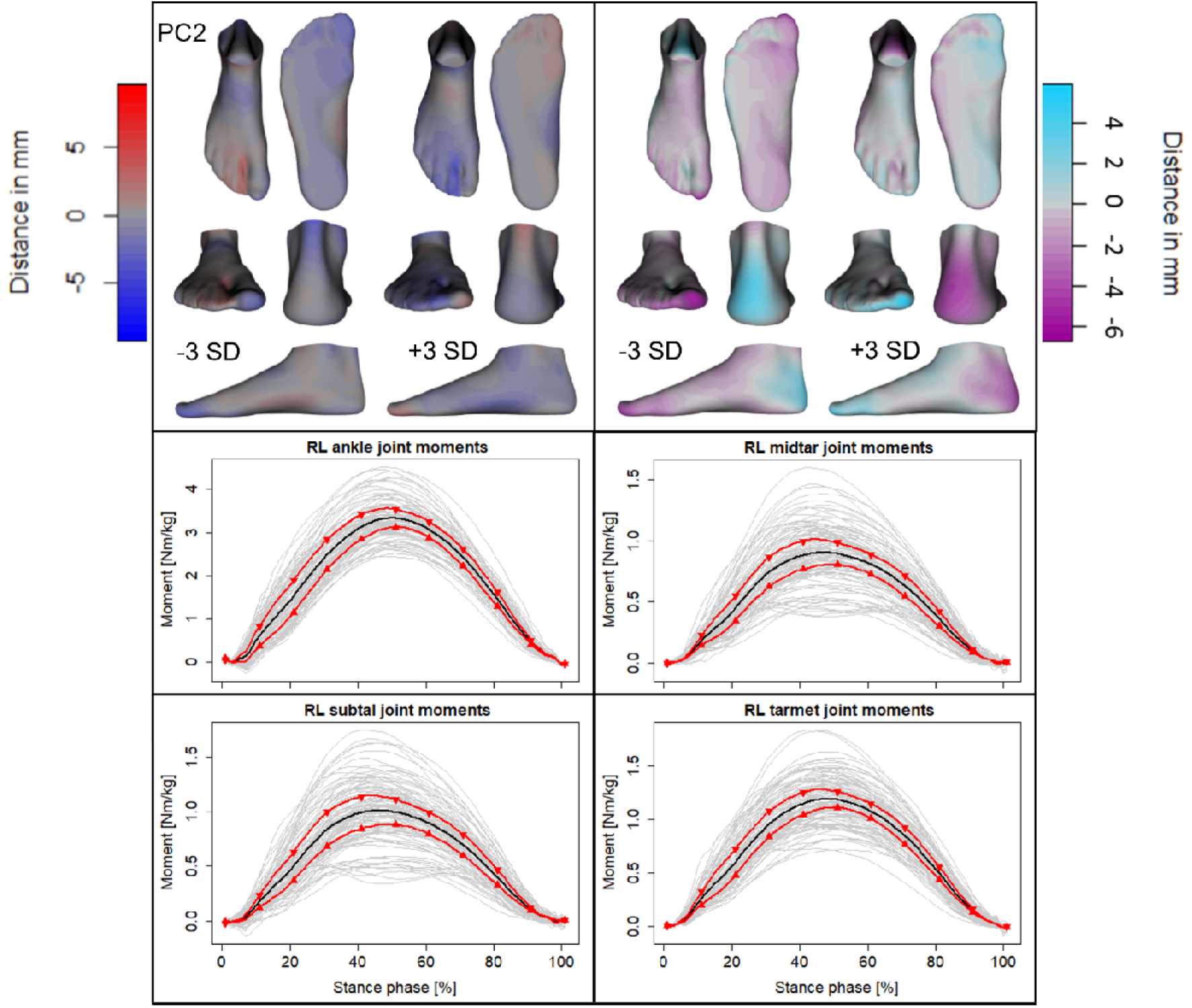
Mean ± 3 standard deviations (SD) of shape (top left), deformation (top right) and moment variations described by the second principal component (PC). Colour coding depicts the degree of deviations from the mean foot or mean deformed foot (Fig. 2). Ankle, subtalar (subtal) and midtarsal (midtar) and tarsometatarsal (tarmet) joint moments during level running (RL) are shown. Upward pointing triangles (▲) denote +3SD and downward pointing triangles (▼) -3SD. Mean (black line) and participant means (grey lines) are also included as a reference.

Accounting for 7.9% of the total variability, the third PC also contained shape differences previously described by statistical foot shape models. In this case, the second PC from these shape models described similar variations in overall foot width and length, TA curvature, and relative length of the lesser toes. The deformations explained by this PC were mainly concerned with the inversion/eversion of both ankle and forefoot and showed a short, wide foot with reduced TA curvature and short toes (−3 SD) to evert and a long, narrow foot with high TA curvature and long toes (+3 SD) to invert at the ankle and forefoot. The former also seemed to exhibit greater subtalar supination and tarsometatarsal plantarflexion throughout stance, along with lower subtalar, midtarsal, tarsometatarsal and MTP joint moments throughout stance, regardless of locomotor task (Fig. 5).

**Fig. 5.**
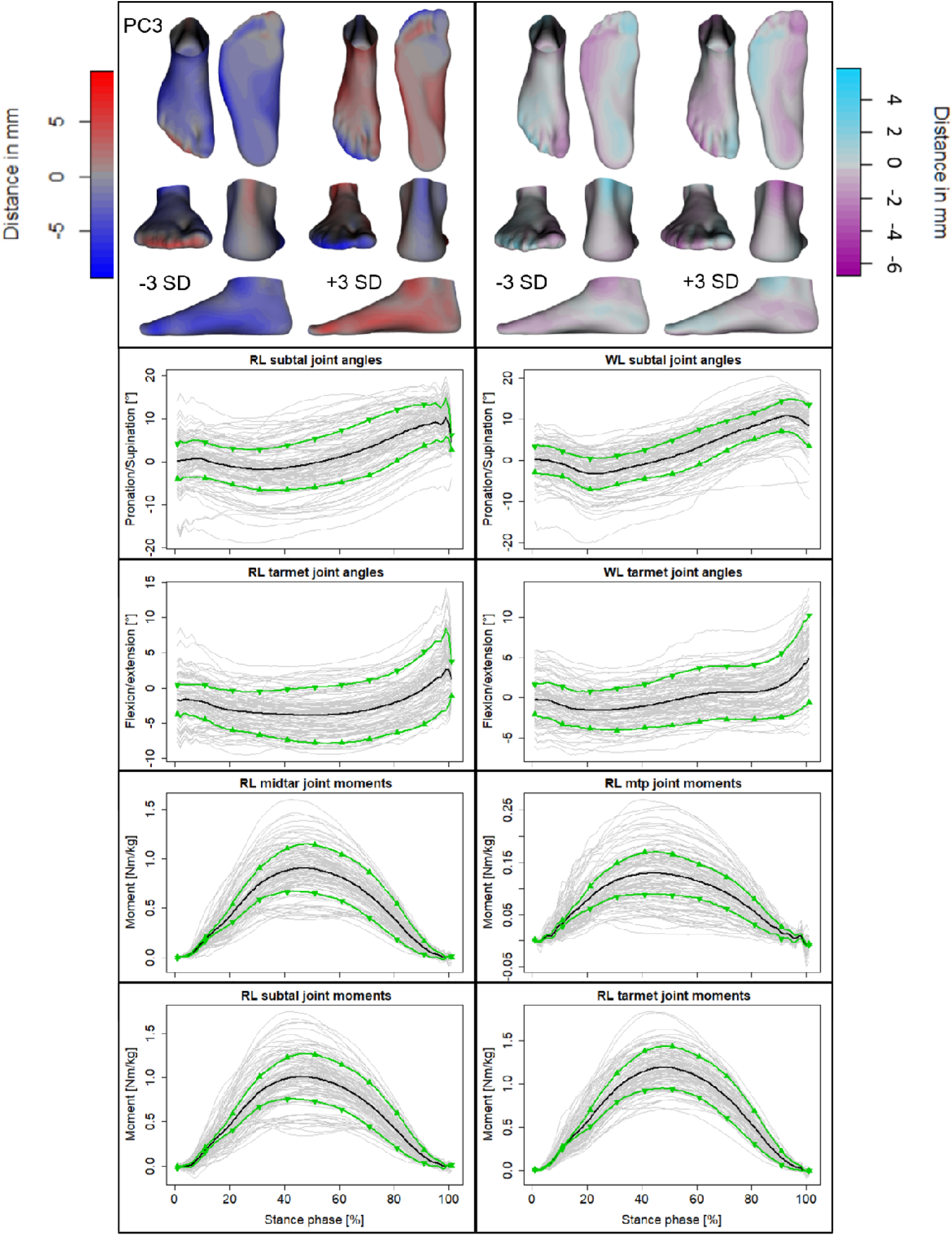
Mean ± 3 standard deviations (SD) of shape (top left), deformation (top right) and joint angle and moment variations described by the third principal component (PC). Colour coding depicts the degree of deviations from the mean foot or mean deformed foot (Fig. 2). Subtalar (subtal) and tarsometatarsal (tarmet) joint angles during level running (RL) and walking (WL) are shown. Joint moments of the midtarsal (midtar), metatarsophalangeal (mtp), subtal and tarmet joints during RL are shown. Upward pointing triangles (▲) denote +3SD and downward pointing triangles (▼) -3SD. Mean (black line) and participant means (grey lines) are also included as a reference.

The 6% variance explained by the fourth PC primarily concerned differences in forefoot, hallux and lesser toe abduction/adduction for both shape and deformation. Already abducted forefoot and toes (−3 SD) was linked to further forefoot and toe abduction when loaded, as well as less tarsometatarsal plantarflexion during late stance (∼80-100%) in running, and early (∼0-20%) and late (∼80-100%) stance in walking. The same foot type was also linked to greater subtalar, tarsometatarsal, midtarsal and MTP joint moments during early stance (∼10-40%) in downhill running, and late stance (∼40-80%) in uphill running (Fig. 6).

**Fig. 6.**
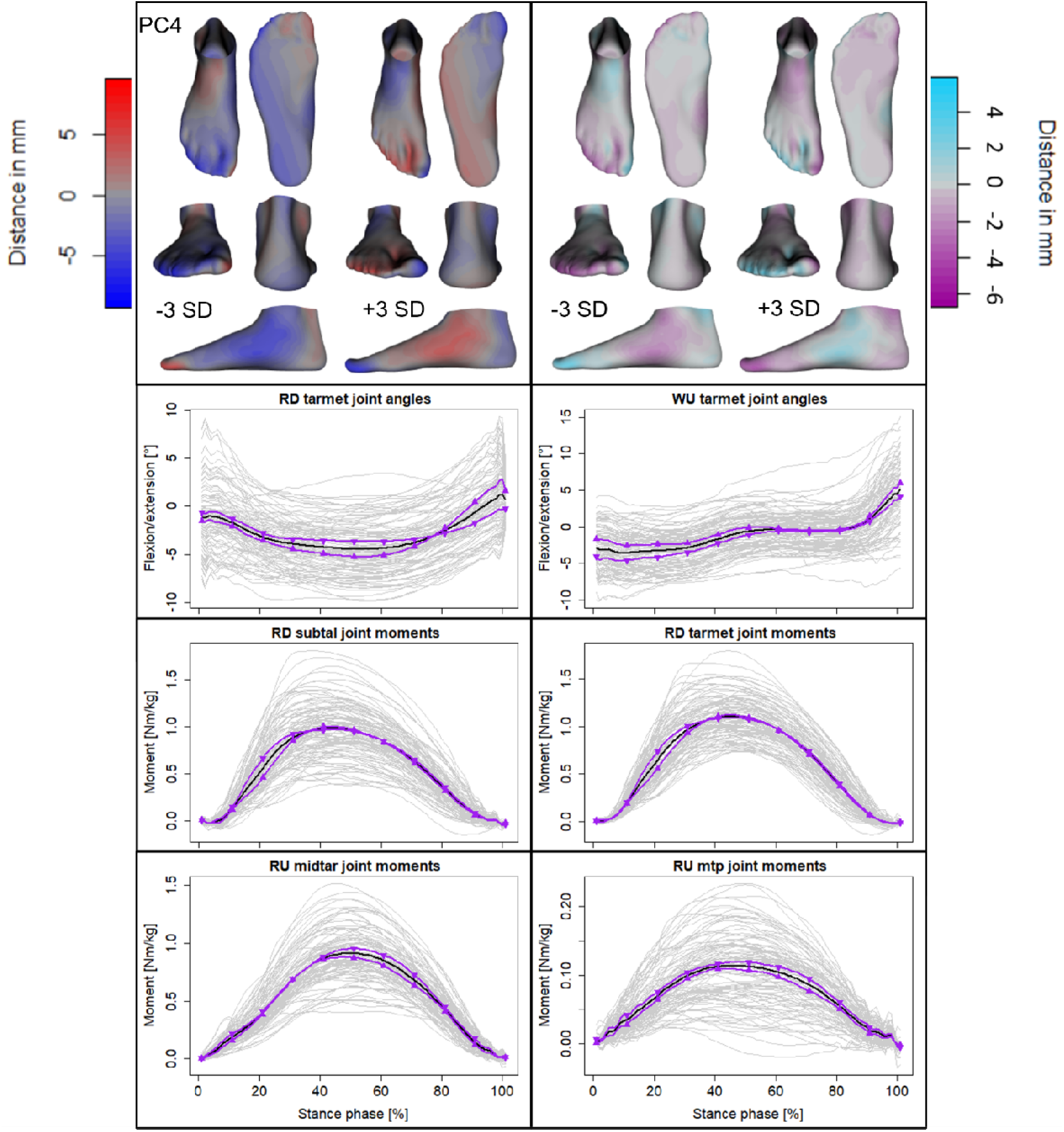
Mean ± 3 standard deviations (SD) of shape (top left), deformation (top right) and joint angle and moment variations described by the fourth principal component (PC). Colour coding depicts the degree of deviations from the mean foot or mean deformed foot (Fig. 2). Tarsometatarsal (tarmet) joint angles during downhill running (RD) and uphill walking (WU) are shown. Joint moments of the subtalar (subtal), tarmet, midtarsal (midtar) and metatarsophalangeal (mtp) joints during RD and uphill running (RU) are shown. Upward pointing triangles (▲) denote +3SD and downward pointing triangles (▼) -3SD. Mean (black line) and participant means (grey lines) are also included as a reference.

The fifth PC only captured 5.6% of the variability in the data. It described shape differences around the height of the ankle, and the separation between the hallux and lesser toes, as well as deformation differences around the splay of the forefoot and toes. Feet characterised by a short ankle and a gap between the hallux and second toe (−3 SD) displayed further separation between hallux and lesser toes, along with lesser toe splay. Furthermore, feet with a tall ankle and no gap between the hallux and second toe (+3 SD) seemed to undergo greater ankle dorsiflexion throughout stance, regardless of locomotor task, while also producing greater MTP joint moments when running uphill (Fig. 7).

**Fig. 7.**
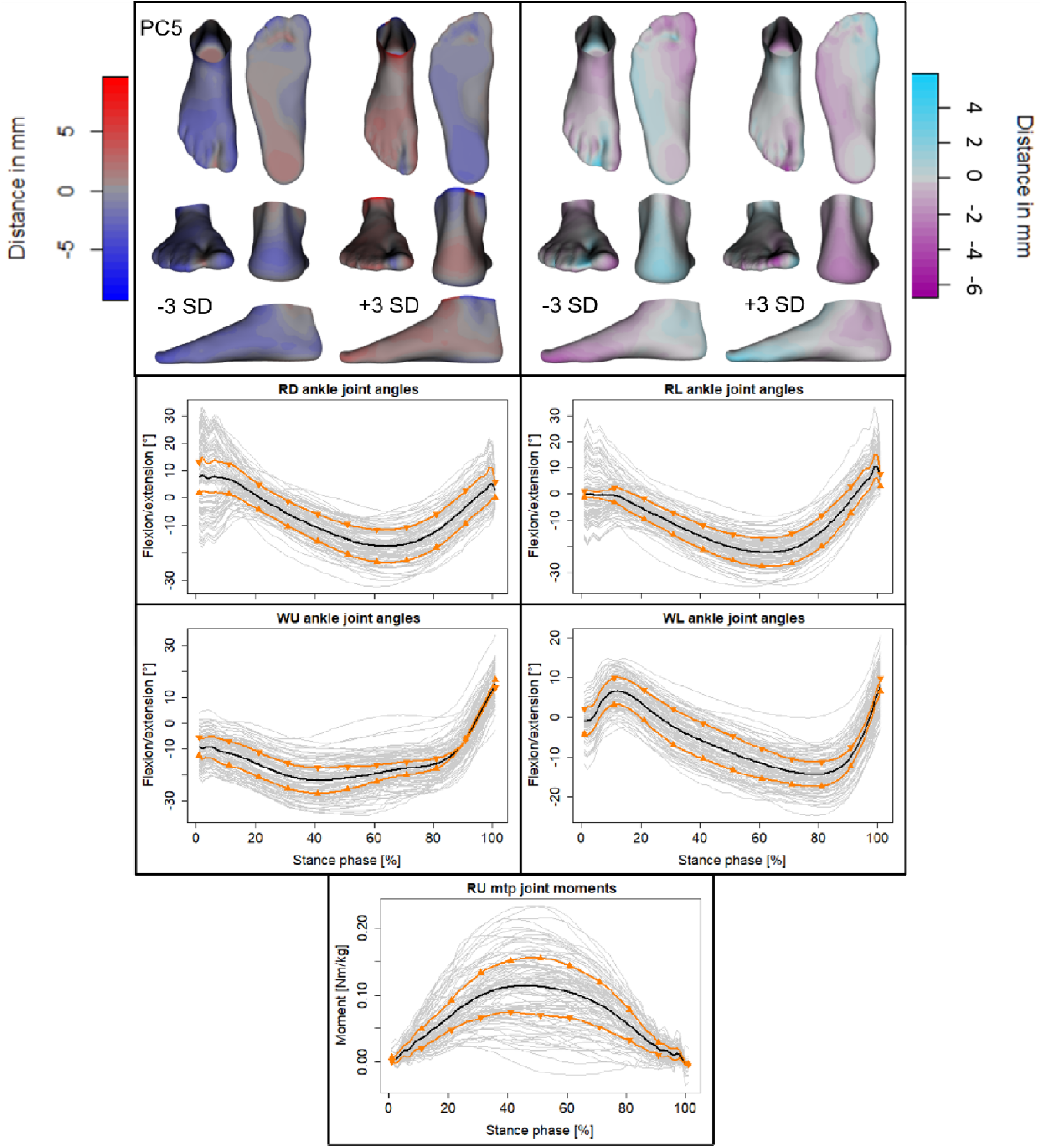
Mean ± 3 standard deviations (SD) of shape (top left), deformation (top right) and joint angle and moment variations described by the fifth principal component (PC). Colour coding depicts the degree of deviations from the mean foot or mean deformed foot (Fig. 2). Ankle joint angles during downhill (RD) and level running (RL) as well as uphill (WU) and level walking (WL) are shown. Joint moments of the metatarsophalangeal (mtp) joints during uphill running (RU) are shown. Upward pointing triangles (▲) denote +3SD and downward pointing triangles (▼) -3SD. Mean (black line) and participant means (grey lines) are also included as a reference.

### Predicting Foot Function from Shape

The LOO analysis, used to assess the accuracy of the SFM when predicting function from shape, produced shape and deformation reconstruction errors that were on average 1.52 ± 1.34 mm and 1.14 ± 1.22 mm, respectively, in magnitude. The mean absolute joint angle prediction errors, averaged across the stance phase, ranged between 1.96 and 5.25° and were generally lowest for the midtarsal and tarsometatarsal joints, as well as the level walking locomotor condition (Fig. 8A). Whereas predictions of ankle and subtalar joint angles, as well as joint angles during uphill and downhill running were least accurate (Fig. 8A). For joint moments, the mean absolute prediction errors ranged between 0.01 and 0.30 Nm/kg. The largest of which were also produced for the ankle joint, while the smallest errors were produced for the MTP joint (Fig. 8C). Similar to the joint angles, level walking produced the lowest errors, while downhill running produced the largest errors across most joints (Fig. 8C).

**Fig. 8.**
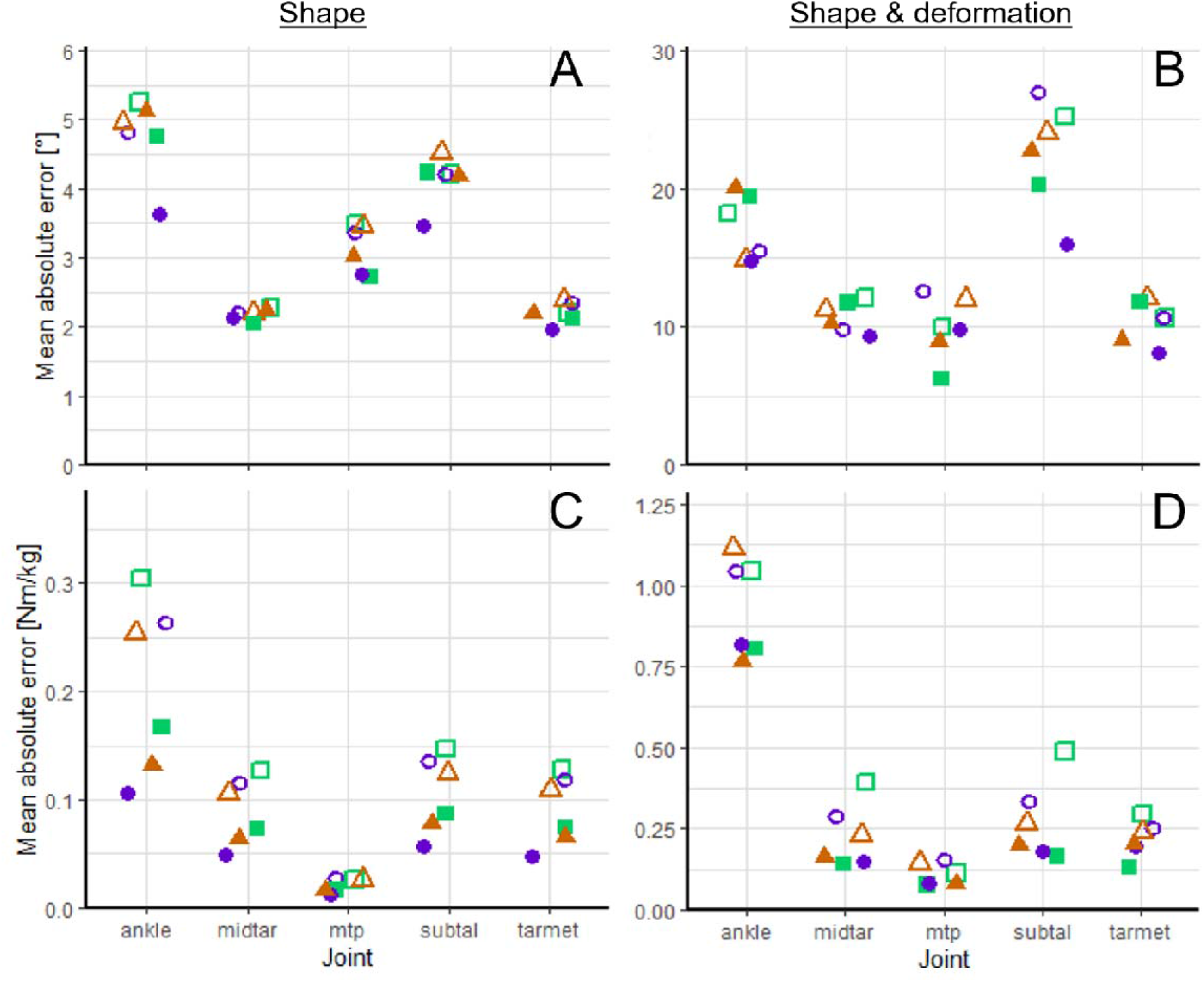
Mean absolute errors between true and predicted joint angles (A and B) and body mass normalised joint moments (C and D). Predictions were made using either just foot shape data (A and C) or foot shape and deformation data (B and D). Results are presented for the ankle, midtarsal (midtar), metatarsophalangeal (mtp), subtalar (subtal) and tarsometatarsal (tarmet) joints, as well as six different locomotor tasks. Noteworthy is that level walking (●) consistently produced small errors and downhill running (□) large errors across all joints. In line with the magnitude of angles and moments experienced at the different joints, midtar and tarmet display small joint angle errors (A and B) and mtp small moment errors (C and D).

Similar patterns regarding the joints and locomotor tasks with the largest and smallest prediction errors were observed for the predicted joint angles and moments when both shape and deformation data was used. However, there was a large difference in magnitude. The mean absolute joint angle prediction errors ranged between 6.21 and 26.96° (Fig. 8B), while joint moment prediction errors ranged between 0.07 and 1.12 Nm kg^-1^ (Fig. 8D). Shape and deformation prediction errors also increased to 7.31 ± 7.36 mm and 5.29 ± 5.05 mm, respectively.

As a reference to compare the errors of predicted foot function against, joint ROM and maximum moments throughout the stance phase are presented in Table 1 as mean ± SD values across locomotor tasks.

**Table 1.**
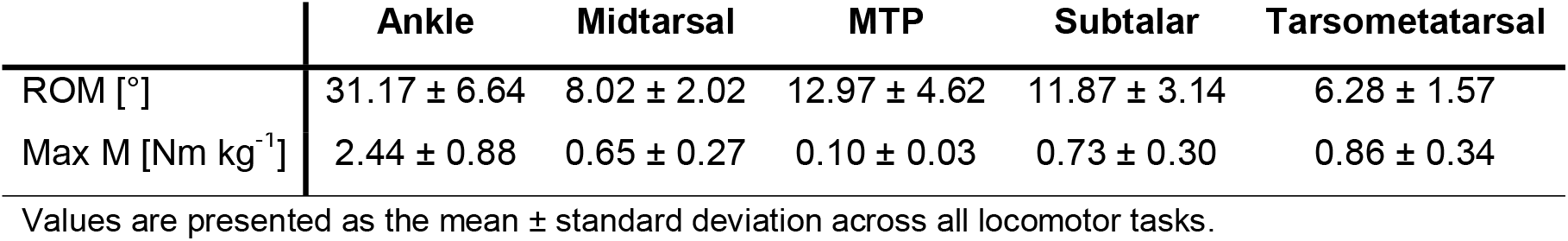
Range of motion (ROM) and maximum body mass normalised joint moment (Max M) for each joint.

## Discussion

The PCA-based statistical shape-function modelling approach taken in this study did not require *a priori* selection of specific predictor and dependent variables. This allows the interpretation of all identified relationships in the context of the foot as a whole, while also avoiding the use of simplified representations of the complex form and function of the foot. An additional advantage of this approach is that the information contained within the SFM can be used to predict the kinematics and kinetics of a new, distinctly shaped foot with reasonable accuracy. Together, the insight provided by this foot SFM and its predictive capacity have the potential to impact a wide range of fields, including evolutionary anthropology, podiatry, orthopaedics and shoe design.

### The Longitudinal and Transverse Arches

Though the results from this investigation support much of the previous research around the LA, they also provide important context relative to the many other components of the foot. The fact that the first PC of the current SFM described LA shape variations, similar to the first PC of prior statistical foot shape models, underscores the magnitude of variation in this aspect of the foot. Moreover, both the variations in load-induced shape deformations, as well as the midtarsal joint kinematics during locomotion also included in this PC support the notion that the LA of a high-arched foot is less likely to compress than that of a low-arched foot, under both static (Arangio et al., 1998; Cornwall and McPoil, 2011; Zifchock et al., 2006; Zifchock et al., 2017) and dynamic loading (Buldt et al., 2015; Kruger et al., 2019; McPoil et al., 2016). Nonetheless, this first PC accounted for only ∼16% of the overall variability, which indicates that the mechanical characteristics of a given foot cannot be defined based solely on the shape of its LA.

Transverse arch curvature covaried with increased midfoot moments and reduced midfoot ROM (PC3), which suggests increased TA curvature is associated with increased LA stiffness. This is the first *in vivo*, locomotion based evidence for the relationship between TA curvature and LA stiffness, as proposed by Venkadesan et al. (2020). Importantly, this finding highlights that foot stiffness is influenced by both LA and TA shape, despite the shapes of the LA and TA appearing to be independent of each other (described in different PCs). This further highlights the multi-factorial nature of foot stiffness and the complex and variable nature of human foot morphology.

The differences in LA shape-associated midtarsal kinematics also seem to translate to kinematic and kinetic differences in the joints proximal and distal to the LA, consistent with the reported mechanical interactions between these joints (Arch and Fylstra, 2016; Carlson et al., 2000; Farris et al., 2020; Magalhães et al., 2021; Smith et al., 2021). Though these mechanical differences appear to exist during level locomotion, they are also apparent during incline and decline conditions. This suggests that differences in LA shape are associated with the mechanical versatility of the foot since the latter locomotor tasks are characterised by either the production of positive (incline locomotion) or negative work (decline locomotion). Furthermore, the involvement of MTP joint mechanics could be an indication that there are differences in how the windlass mechanism affects the mechanical behaviour of feet with differently shaped LAs, forefeet and heels. However, in the absence of electromyographic data, whether these mechanical differences are due to differences in active muscular control (Caravaggi et al., 2010; Farris et al., 2020; Farris et al., 2019; Kelly et al., 2019; Kelly et al., 2015) or passive action (Erdemir et al., 2004; Hicks, 1954; Welte et al., 2021; Welte et al., 2018), or their combination, cannot be defined and would require further investigation.

### Relative Foot Proportions

Another noteworthy finding is that ankle, subtalar, midtarsal, tarsometatarsal and MTP joint moments are associated with the relative proportions of the foot, as described in PCs 2 and 3. In the latter, larger joint moments coincide with a long and narrow foot shape (+3 SD). This link can be explained by the fact that as relative foot width decreases and relative foot length increases, so does the moment arm between the respective GRF components’ point of application and the joints’ centre of rotation. However, in PC2 it is a relatively longer forefoot and shorter heel (−3 SD) that are linked to larger ankle, subtalar and midfoot joint moments. Specifically the larger ankle moments despite similar ROM seem analogous to previous reports of increased plantar flexor work as a consequence of an increased forefoot to rearfoot length ratio (Baxter et al., 2012), increased elastic energy reutilisation in the Achilles tendon associated with a shorter calcaneal tuber length (Raichlen et al., 2011), and increased elastic energy reutilisation in the ankle spring of a prothesis with longer LA length (Honert et al., 2020).

Interestingly, the variations in foot proportions described by PC2 are only associated with mechanical differences during level running, whereas those described by PC3 are associated with mechanical differences across all locomotor tasks. This task dependence could be due to the specific energetics of each task. The previously mentioned suggestion of an increase in Achilles tendon elastic energy reutilization due to a shorter heel was found to affect running but not walking economy (Raichlen et al., 2011). Similarly, the elastic energy stored and returned through compression and recoil of the LA was found to reduce the metabolic cost only of level running but not walking nor uphill running (Stearne et al., 2016). Moreover, the metabolic cost of level running in general, unlike walking, is heavily reliant on elastic recoil (Alexander, 1991). Thus, the bouncing spring-mass, which level running has been likened to (Blickhan, 1989), may be the only condition that produces the ankle and midfoot moments beneficial for elastic energy storage, especially in feet with the relative proportions described by PC2. On the other hand, greater joint moments due to increased moment arms, as in PC3, will occur regardless of locomotor task. Further insight into these form-function associations could be also provided by information regarding the muscular activation patterns during these different locomotor tasks, as well as their associations to the specific foot shape characteristics.

### Hallux and Lesser Toe Lengths and Orientation

Further notable differences between PCs 2 and 3 lie in the variations in toe length and MTP joint moments. In PC2 only the hallux varies in length, whereas in PC3 it is the length of the lesser toes that varies. While some investigations have found that neither orientation nor length (absolute or relative) of the toes affect walking dynamics (Honert et al., 2018; Honert et al., 2020), others have found that toe length does affect running performance (Rolian et al., 2009; Tanaka et al., 2017; Ueno et al., 2018). As proposed by the latter two investigations, longer toes (especially a longer second toe) create larger MTP joint moments, which in turn affects plantar flexor gearing in such a way that is beneficial to both sprinting (Tanaka et al., 2017) and endurance running (Ueno et al., 2018). Correspondingly, in our SFM, variations in hallux length are associated with only small variations in MTP joint angles and moments (PC2), whereas variations in lesser toe lengths are associated with large variations in MTP joint angles and moments (PC3).

Additional shape characteristics of the toes, such as their orientation (PC4) and the separation between hallux and lesser toes (PC5), are also associated with variations in MTP mechanics at specific points during the stance phase, especially during uphill and downhill running. However, regardless of the PC, toe shape characteristics are not the only variations in foot shape described and therefore cannot be interpreted as the only source of mechanical variations at the MTP joint. The evidence for a mechanical connection between the ankle and foot joints, as discussed for PC1, indicates that forefoot abduction (PC4) and ankle height (PC5) may also be partly responsible for the mechanical differences seen at the MTP joint in these PCs. Nevertheless, the inclusion of some aspect of toe shape in each of the first five PCs demonstrates the large variability herein while also aligning with some previous studies that have examined the mechanical implications of varying toe shape characteristics.

### Predicting Foot Function from Shape

The information about the external shape of a given foot can be used to predict its mechanical function. When only shape is used to predict function, the errors are generally within the standard deviations in mean ROM and maximum joint moments across locomotor tasks (Table 1). However, including deformation data as a predictor of mechanical function, results in drastically reduced accuracy. This could be because the shape deformations under static loading are not as closely related to locomotor function (Farris et al., 2020; Sichting and Ebrecht, 2021; Welte et al., 2018; Williams et al., 2022) as shape alone appears to be. Additionally, the added inter-subject variability and potential noise from unloaded foot shape contained in the deformation data (Schuster et al., 2021) may also contribute to the reduced predictive accuracy.

### General Implications of the Statistical Shape-Function Model

Above all else, the results from this investigation highlight the complexity and high variability of the foot, both in form and function. The largest PC, as well as the following five required to explain just over half of the total variability, all account for relatively small proportions of the total variability (i.e., small eigenvalues). This shows that, due to its structural complexity, the relationships between foot form and function may not be tightly constrained. In comparison, similar SFMs of less complex biological structures, such as the knee, have accounted for over 45% of the variability in shape and function with just the first PC (Fitzpatrick et al., 2011a; Smoger et al., 2015). Moreover, the small relative weight of the specific shape-function relationships described by each PC to overall shape-function variation is comparable to previous reports of weak-to-moderate relationships between the shape and function characteristics of specific foot structures (Cavanagh et al., 1997; Nielsen et al., 2010; Nigg et al., 1993; Zifchock et al., 2006). However, our SFM is the first to simultaneously consider all of the previously described shape-function relationships and assign them a hierarchical order of importance. Thereby confirming the relative significance and large variability of the LA, while also emphasising the importance of foot proportions, and toe orientation and length.

The small eigenvalues also imply that the large degrees of freedom afforded by the structural complexity of the foot allow for multiple means of achieving the same functional outcome. That is, despite large variations in foot shape and mechanics, all participants were able to successfully perform each of the locomotor tasks. As Lauder (1995) argues, function is not defined by musculoskeletal structure itself, but by how it is used, i.e., “the motor programs in the central nervous system”. There is ample evidence for this argument in relation to the foot, as various investigations have shown that the neuromuscular control of intrinsic and extrinsic foot muscles plays an important role in the mechanical function and versatility of the foot (Birch et al., 2021; Kessler et al., 2020; Riddick et al., 2019; Smith et al., 2021). Thus, future investigations examining the link between foot form and function might benefit from also examining the potential link to the neuromuscular control of relevant muscles during various locomotor tasks. Furthermore, Lauder (1995) points out that the definition of function can vary widely. In the current investigation it was defined as the joint kinematics and kinetics produced to complete locomotor tasks with varying mechanical requirements. However, many other investigations have defined function as the metabolic cost of locomotion (Charles et al., 2021; Raichlen et al., 2011; Stearne et al., 2016) or performance during a locomotor task (Tanaka et al., 2017; Ueno et al., 2018). Since these definitions of function have greater implications for general human behaviour in an evolutionary context, linking foot form and mechanics to locomotor economy using a similar approach as that used here might help resolve some of the remaining questions regarding the link between foot structure and locomotor economy, such as whether there are also variations in anatomical parameters that are as closely related to walking economy (Charles et al., 2021) as heel length is to running economy (Raichlen et al., 2011), or whether short (Rolian et al., 2009) or long toes (Ueno et al., 2018) are better for endurance running?

The information around the range of variation in foot form and function, the relationships between these two, and the predictive capacity of the SFM all have applications across a broad range of fields. The wide variability in both foot shape and function contained within the SFM could be used as a comparison against the proposed shape and function of fossilised feet and footprints for a better understanding of how far modern human foot function diverges from that of our evolutionary ancestors. The various form-function relationships described by the SFM could also be used to inform the design of shoes aimed at enhancing running performance, since the impact shoe design can have on performance has been found to vary considerably across runners (Hébert-Losier et al., 2022). Based on the form-function relationships discussed above, a low-arched foot might benefit more from increased midsole bending stiffness, whereas a foot with a relatively long heel and short forefoot might benefit more from a compliant midsole. Thus, it might be possible to determine the ideal shoe for a given runner based on their foot shape characteristics. Lastly, the predictive capacity of the SFM could provide a more complete understanding of the inferred mechanical function of fossilised feet and footprints if their complete external shape can be reconstructed from partial data, as has been done for skeletal foot segments (Grant et al., 2020). Similarly, comparing the predicted healthy mechanics to those measured from an impaired foot could be used as a diagnostic tool by podiatrists and orthopaedic surgeons.

### Limitations

The findings of this study must be considered in light of some limitations. Firstly, a limitation that is common to studies using a PCA-based approach to identify the major modes of variation or covariation in a set of complex data is that the interpretation of the resulting PCs can be somewhat subjective. To help reduce subjectivity in the current interpretation of foot shape and deformation, the differences therein were highlighted using heatmaps that denote the magnitude of deviation from the mean. Animations (included in the supplementary information) were also used to further emphasise the areas and nature of shape changes. An additional approach to further reduce subjectivity in future investigations could be to correlate common measures of shape characteristics (e.g., foot length, width, arch height or arch index obtained from the 3D foot scans) to their respective PC scores, as Smoger et al. (2015) have done. However, such an approach was outside of the scope of the current study. In relation to the foot function data, a limitation to consider is that only the kinematics and kinetics during the stance phase were analysed. Differences during the swing phase might add information that could help further refine the currently observed shape-function relationships. Additional factors such as spatiotemporal gait variables (stride length and frequency) might also account for some of the variation in kinematics and kinetics. Lastly, considering that our sample consisted of only young, healthy participants, the results presented here may not be transferable to clinical, infant or elderly populations.

### Conclusion

Using a PCA-based approach to identify relationships between external foot shape, deformations and joint mechanics during locomotion has revealed that, across a healthy, young population, the greatest variability exists in the LA, TA, relative foot proportions and toe shape characteristics as well as their associated deformations and joint mechanics. While the relationships between these various aspects of external foot shape and internal joint mechanics agree with similar findings from previous research, context regarding their relevance to overall foot form and function is provided for the first time by the relatively small proportion of overall variability accounted for by each of these covariations. Beyond describing the various links between shape characteristics and joint mechanics, the foot SFM presented here has potential applicability across a wide range of fields, since it can be used to predict, with reasonable accuracy, the internal joint mechanics of a given foot based only on its external shape.

## List of Abbreviations and Symbols

3D: Three-dimensional
fBW: Full body weight
GRF: Ground reaction force
LA: Longitudinal arch
LOO: Leave-one-out
mBW: Minimal weight
MTP: Metatarsophalangeal
PC: principal component
PCA: Principal component analysis
SD: Standard deviation
SFM: Shape-function model
TA: Transverse arch

## Acknowledgements

The authors would like to thank Max Andrews for his help collecting the data. The authors would also like to thank ASICS Oceania for providing complementary footwear to all participants.

## Competing interests

No competing interests declared.

## Author contributions

R.W.S. and L.A.K. conceived and designed the experiments. R.W.S. collected, processed and analysed the data. All authors discussed the results and contributed to the elaboration of the manuscript. All authors have read and approved the final version of this manuscript.

## Funding

This research was funded by an Australian Research Council (ARC) Discovery Early Career Research Award [DE200100585 to L.A.K.].

## Data availability

The 3D foot scans (PLY files), and locomotor foot and ankle joint angles and moments (MatLab structure) can be downloaded from the UQ Research Data Manager:

https://doi.org/10.48610/4a4dc49

## Notes

### Competing Interest Statement

The authors have declared no competing interest.

